# Evolution of herbivory remodels a *Drosophila* genome

**DOI:** 10.1101/767160

**Authors:** Andrew D. Gloss, Anna C. Nelson Dittrich, Richard T. Lapoint, Benjamin Goldman-Huertas, Kirsten I. Verster, Julianne L. Pelaez, Andrew D. L. Nelson, Jessica Aguilar, Ellie Armstrong, Joseph L.M. Charboneau, Simon C. Groen, David H. Hembry, Christopher J. Ochoa, Timothy K. O’Connor, Stefan Prost, Hiromu C. Suzuki, Sophie Zaaijer, Paul. D. Nabity, Noah K. Whiteman

## Abstract

One-quarter of extant Eukaryotic species are herbivorous insects, yet the genomic basis of this extraordinary adaptive radiation is unclear. Recently-derived herbivorous species hold promise for understanding how colonization of living plant tissues shaped the evolution of herbivore genomes. Here, we characterized exceptional patterns of evolution coupled with a recent (<15 mya) transition to herbivory of mustard plants (Brassicaceae, including *Arabidopsis thaliana*) in the fly genus *Scaptomyza,* nested within the paraphyletic genus *Drosophila*. We discovered a radiation of mustard-specialized *Scaptomyza* species, comparable in diversity to the *Drosophila melanogaster* species subgroup. Stable isotope, behavioral, and viability assays revealed these flies are obligate herbivores. Genome sequencing of one species, *S. flava,* revealed that the evolution of herbivory drove a contraction in gene families involved in chemosensation and xenobiotic metabolism. Against this backdrop of losses, highly targeted gains (“blooms”) were found in Phase I and Phase II detoxification gene sub-families, including glutathione *S*-transferase (*Gst*) and cytochrome P450 (*Cyp450*) genes. *S. flava* has more validated paralogs of a single *Cyp450* (N=6 for *Cyp6g1*) and *Gst* (N=5 for *GstE5-8*) than any other drosophilid. Functional studies of the *Gst* repertoire in *S. flava* showed that transcription of *S. flava GstE5-8* paralogs was differentially regulated by dietary mustard oils, and of 22 heterologously expressed cytosolic *S. flava* GST enzymes, GSTE5-8 enzymes were exceptionally well-adapted to mustard oil detoxification *in vitro.* One, GSTE5-8a, was an order of magnitude more efficient at metabolizing mustard oils than GSTs from any other metazoan. The serendipitous intersection of two genetic model organisms, *Drosophila* and *Arabidopsis,* helped illuminate how an insect genome was remodeled during the evolutionary transformation to herbivory, identifying mechanisms that facilitated the evolution of the most diverse guild of animal life.

**SIGNIFICANCE STATEMENT:** The origin of land plants >400 million years ago (mya) spurred the diversification of plant-feeding (herbivorous) insects and triggered an ongoing chemical co-evolutionary arms race. Because ancestors of most herbivorous insects first colonized plants >200 mya, the sands of time have buried evidence of how their genomes changed with their diet. We leveraged the serendipitous intersection of two genetic model systems: a close relative of yeast-feeding fruit fly (*Drosophila melanogaste*r), the “wasabi fly” (*Scaptomyza flava*), that evolved to consume mustard plants including *Arabidopsis thaliana*. The yeast-to-mustard dietary transition remodeled the fly’s gene repertoire for sensing and detoxifying chemicals. Although many genes were lost, some underwent duplications that encode the most efficient detoxifying enzymes against mustard oils known from animals.

## INTRODUCTION

The origin of land plants over 400 million years ago presented a new niche for insects (1), ultimately producing the most diverse assemblages of animal life ever to have evolved (2, 3). Herbivory explains ~25% or more of the variation in diversification rates and species richness across insect orders (4, 5), and has resulted in convergent evolution across many levels of biological organization, from morphology and behavior to genes (6) and even single amino acids (7–9).

An unresolved problem is whether repeated herbivore diversification events spur concomitant diversification of gene families with functions intimately involved in the interaction between insects and plants. Of particular interest are genes involved in mediating interactions with plant secondary compounds, which include toxins that must be mitigated, yet are also often used as host-finding cues by specialized herbivores. Escalating chemical co-evolutionary arms races between plant defense and herbivore counter-defense are the preeminent mechanism hypothesized to drive the origin of new species, adaptations, and genes, on both sides of the equation (10–14).

Comparative genomics offers a powerful lens for uncovering the adaptations facilitating herbivory. For example, the spider mite (*Tetranychus urticae*) and diamondback moth (*Plutella xylostella*) genomes harbor expanded gene families encoding enzymes involved in detoxification of plant secondary compounds (15–17), and genes encoding gustatory receptors (GRs) involved in host finding have undergone extensive lineage-specific duplications in butterflies (18). Although there are clear gene-family expansions arising from plant-insect interactions, including “blooms” of paralogs that emerge from single ancestral genes (13, 19), isolating herbivory as the causal agent of some of these exceptional genome-scale patterns remains controversial (20). Herbivory evolved >25 million years ago in all of the eighteen arthropod lineages assayed using comparative genomics (Fig. 1a). The most diverse living lineages, which include butterflies and moths (Lepidoptera), and leaf beetles, weevils and close relatives (Phytophaga), arose in the early Mesozoic (5). The lack of more closely related non-herbivorous lineages for comparison makes it difficult to pinpoint the timing of these genomic changes relative to the evolution of herbivory, and thus to distinguish herbivory-associated patterns from those that might have arisen in response to other factors or even through stochastic processes. Further, these patterns may reflect subsequent evolutionary consequences of herbivory, rather than the mechanisms facilitating the evolution of herbivory in the first place. Thus, while clear patterns have been uncovered in the genomes of some herbivorous insects (21), establishing that herbivory drove these genomic patterns is a challenge, just as disentangling cause and effect is a principal challenge facing comparative evolutionary genomics studies generally.

**Figure 1.**
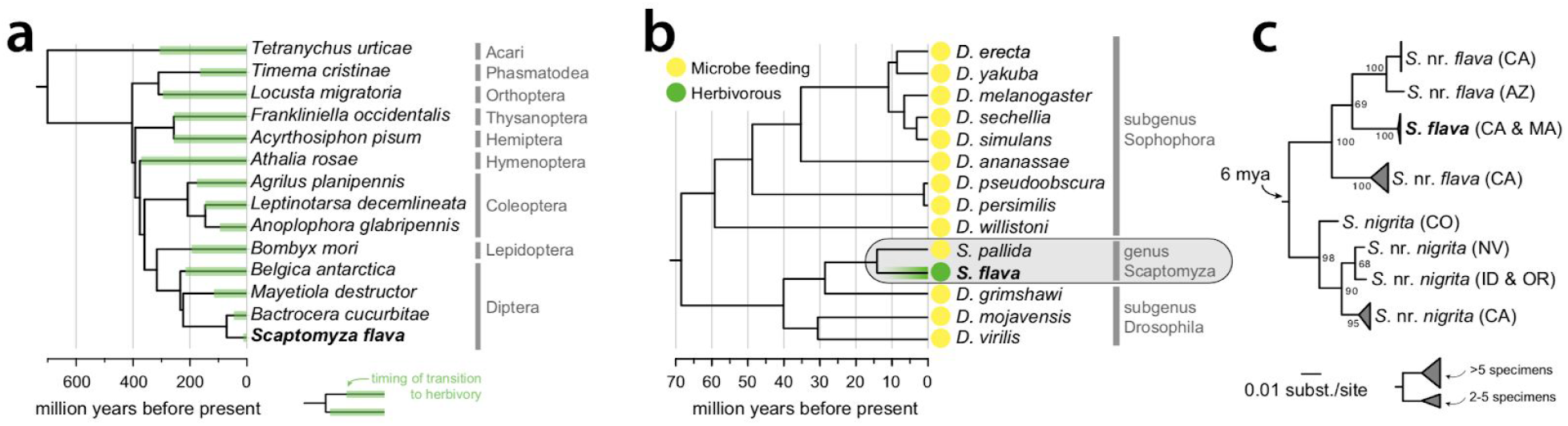
A young herbivorous lineage in the Drosophilidae. **(a)** Likely timing of transitions to herbivory, inferred from the reported diet breadth of related families and orders, in the ancestors of terrestrial herbivorous arthropods with sequenced genomes, **(b)** Phylogenetic placement of *S. flava* within the paraphyletic genus *Drosophila (23*). Shown are the first 12 *Drosophila* species to have their genomes sequenced, along with the closest known non-herbivorous relative of *S. flava* to clarify the timing of the transition to herbivory (30). **(c)** Maximum likelihood nucleotide phylogeny built using *COI* sequences from North American *Scaptomyza* collected feeding on mustard plants (Brassicales spp.). Individuals separated by < 1% pairwise nucleotide divergence were iteratively collapsed, such that clades are distinct at a 1% divergence cutoff. Collection localities are indicated by two-letter state abbreviations.

One potential advance is to use comparisons among more closely related (recently diverged) taxa. Although genomic changes may be subtler when relatively short intervals have elapsed since herbivory evolved, comparing young herbivorous lineages with many closely related non-herbivorous sister taxa offer a number of advantages. This approach can identify genomic patterns in herbivores that are unmatched across non-herbivorous relatives, more precisely pinpoint the timing of these changes, and uncover the specific genes driving these patterns as well as other molecular evolutionary details.

True flies (Diptera) are attractive as models to address these questions: herbivory has evolved at least 25 times independently and many times relatively recently (22). Within Diptera, the family Drosophilidae (including the genus *Drosophila*) presents an exceptional opportunity to dissect the molecular basis of adaptations facilitating the evolution of herbivory. Although most species within Drosophilidae retain the ancestral habit of feeding on microbes associated with decaying plant tissue, herbivory has evolved several times in the past 20 million years (23). An in-depth understanding of the functions of thousands of genes in *Drosophila,* coupled with the availability of high quality genome sequences from species spanning the phylogeny, offers resources unparalleled in other insect lineages (24).

Notably, one of these transitions to herbivory involved colonization of plants in the Brassicales, including the genetic model plant *Arabidopsis thaliana* (26). This transition occurred within the genus *Scaptomyza* (Fig. 1b), which is nested phylogenetically within subgenus *Drosophila* (sister to Hawaiian *Drosophila*? (25)). Despite little attention from taxonomists, molecular barcoding suggests this transition is associated with a cryptic species radiation (8 species in North America alone) similar in species richness and phylogenetic diversity to that found in the *D. melanogaster* subgroup worldwide (9 species) (Fig. 1c) (27). To probe the evolutionary genomics of this recent transition to herbivory, we merged pre-existing genetic and genomic resources for *Drosophila* and *Arabidopsis* with new genome sequences from an herbivorous *Scaptomyza* species (*S. flava*). These resources have enabled detailed evolutionary and functional characterization of genomic patterns associated with herbivory.

Our analyses revealed a slight overall contraction in chemosensory and detoxification gene number. This pattern belied elevated, partially-offsetting rates of gene gains and losses, with “blooms” of genes that function in the interaction with dietary or environmental compounds and toxins. Expression of five *Scaptomyza-*specific paralogs of *GstE5-8,* the product of one such bloom, were highly regulated by dietary mustard oils, the primary defensive toxins in its host plants. Michaelis-Menton kinetics showed that three of these GSTs were the most efficient at detoxifying mustard oils of any known from animals. We also found six paralogous copies of the CYP450 gene *Cyp6g1,* which in several other drosophilids has experienced recent positive selection for synthetic insecticide resistance through increases in copy number and gene expression (28, 29). Together, our analyses uncover dynamic gene family remodeling in an herbivore, illuminated by comparisons with a suite of relatives that retained an ancestral, non-herbivorous feeding mode.

## RESULTS & DISCUSSION

### An obligately herbivorous drosophilid

The paraphyletic genus *Drosophila,* including the genus *Scaptomyza* nested within it, is composed of >2,000 species that have diversified across a wide range of feeding niches (31). While many drosophilids feed on rotting plant or fungal tissue and their associated microbes, including a majority of species in the subgenus *Drosophila* and the genus *Scaptomyza* (25, 31), only a handful of species are reported to feed on living plants (24). We first explored the trophic level occupied by *S. flava* (Fig. 2a) using behavioral assays, viability assays, and stable isotope profiling to determine if it truly consumes and derives nutrients from living plant tissue.

**Figure 2.**
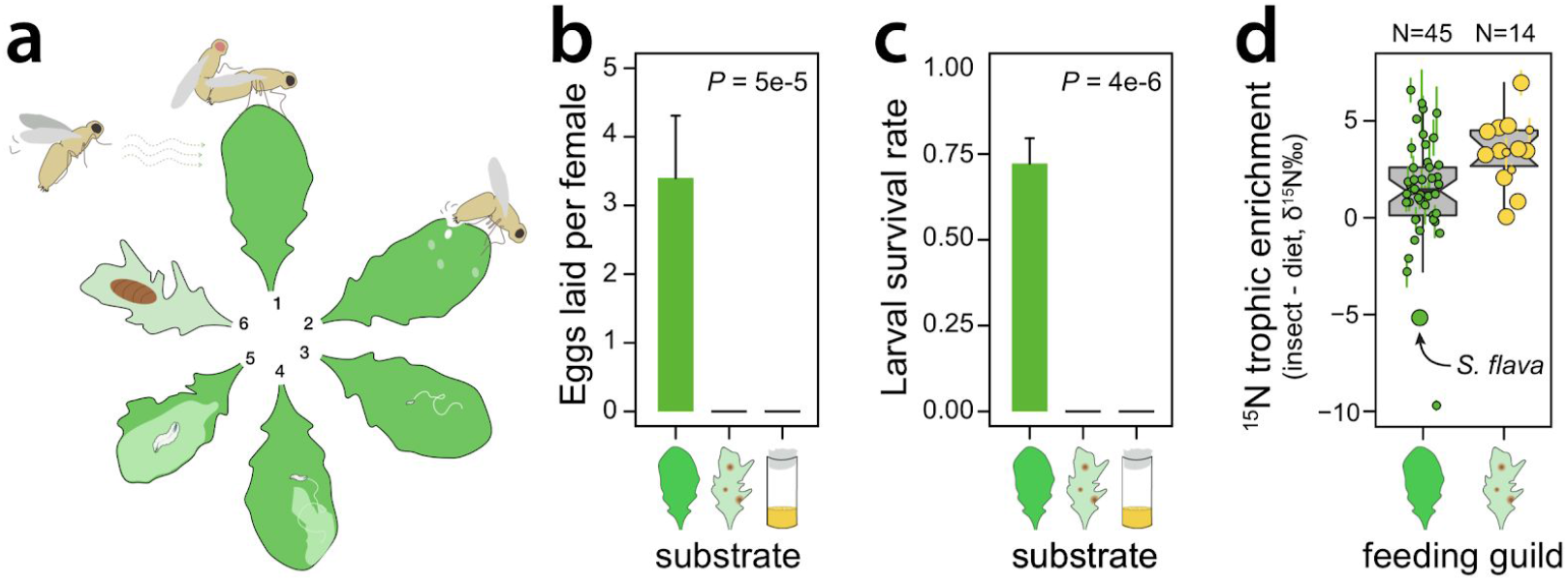
*S. flava* is an obligate herbivore, **(a)** The life cycle of *S. flava* is intimately linked with its host plants: (1) adults mate on or near host plants; (2) adult females pierce plants with their serrated ovipositor, feeding on the exudates and depositing eggs into the wounds; (3–5) larvae progress through three instars as leafmining endoparasites; and (6) pupation occurs on or near host plants, **(b)** Adult female *S. flava* exclusively oviposit in living plant tissue in three-way choice tests between living Col-0 plants, rotting Col-0 plants, and yeast-seeded *Drosophila* growth media (Kruskal-Wallis test; N = 34 total eggs across 10 trials), **(c)** Following transfers to the same substrates, larval *S. flava* only survive to adulthood on living Col-0 plants (Kruskal-Wallis test; N = 9 replicates per substrate, with 6 larvae per replicate), **(d)** Trophic enrichment of heavy nitrogen isotopes (^15^N) in insects relative to their feeding substrate, for insects consuming living vs. decaying, microbe-rich plant material. Large circles indicate observations from *S. flava* or microbe-feeding *Drosophila* (this study and previous studies), while small circles indicate observations from other insects (previous studies; Table S1). Each circle indicates an averaged observation for a combination of insect species and host species/substrate +/− SEM. Circles are offset on the x-axis to aid in visualization

We censused eggs laid by *S. flava* in a three-way choice test offering equal proportions of living *A. thaliana* Col-0 plants, rotting Col-0 plants, and yeast (Fleischmann’s)-seeded nutrient media. Eggs were deposited exclusively in the leaves of living plants (Fig. 2b), consistent with the observation that adult *S. flava* antennae are responsive to volatiles from green plants but not from yeast in olfactory assays (32). Next, we transferred second instar larvae onto each of the same three substrates. Larvae only completed development to adults in living plant tissue (Fig. 2c). Thus, *S. flava* is an obligate herbivore.

These assays, however, do not necessarily demonstrate that *S. flava* obtains nutrients directly from plant tissue. An alternative possibility is that *S. flava* consumes a mixture of living plant tissue and associated microbes, but only digests and obtains nutrients from the latter. Comparisons of stable isotope composition between insects and putative dietary substrates offer insight into nutrient acquisition: lighter isotopes are lost preferentially relative to heavier ones as nutrients move up trophic levels, leading to an enrichment of heavy nitrogen isotopes (^15^N). We profiled nitrogen isotope composition in *S. flava* and its Col-0 host plants and in *Drosophila* and their rotting fruit substrates, and then compared these to additional profiles from the literature. As expected, *Drosophila* and other insects feeding on rotting, microbe-rich substrates typically exhibited a two-fold greater trophic enrichment of ^15^N than herbivores (Fig. 2d), reflecting the additional trophic step between insect and dietary substrate (i.e., rotting plant tissue → microbe → insect, vs. living plant → insect; (33)). In contrast, ^15^N was depleted in *S. flava* relative to its host plants (Fig. 2d). Trophic depletion of ^15^N has been observed in fluid-feeding herbivorous insects (34, 35), but it is inconsistent with a diet consisting primarily of plant-associated microbes. Our finding that *S. flava* primarily derives nutrients from living plant tissue is further supported by previous findings that larval feeding and growth rates are not directly enhanced by elevated bacterial loads in their diet, although larvae do benefit indirectly through bacterial suppression of plant defenses (36). Overall, these experiments show that *S. flava* is an obligate herbivore that consumes and acquires nutrients from living plant tissue.

### The *S. flava* genome

Next, we proceeded to examine the genomic consequences of this transition to herbivory. To enable comparative genomic studies, we sequenced, assembled, and annotated the genome of a fully inbred *S. flava* line collected as larvae from *Barbarea vulgaris* at Beaver Brook Reservation in Belmont, MA, USA in 2008. A total of 26 Gb of paired-end Illumina reads (approximately 90x coverage of the estimated 289 Mb genome; (26)), from libraries with insert sizes ranging from 180 bp to 5 kb, produced a 203 Mb assembly with a scaffold N50 size of 118 kb. We further scaffolded and extended contigs using PacBio reads, increasing the N50 to 211 kb. Using gene models from other *Drosophila* and RNA sequenced across *S. flava* life stages, we predicted 12,993 and 17,997 genes using high confidence and liberal methods, respectively. The gene space of the assembly is near complete: only 1.5% of 2799 core dipteran genes are absent, which falls near the median value for the 12 species of the original Sanger sequenced genome assemblies that form the foundation for *Drosophila* comparative genomics (Tables S2-S4; (30)).

To validate genes and genomic regions of interest, we also leveraged a separate *S. flava* genome assembly generated from long-read sequencing and Hi-C scaffolding of a partially inbred *S. flava* colony collected near Dover, NH, USA. This assembly has a higher number of missing genes than the primary assembly, but also offers much longer contiguous stretches that are informative for distinguishing duplicated genes from separately-assembled alleles (Tables S2-S3). Preparation for the release of this assembly, which is available upon request, is ongoing.

### Contraction and turnover of chemosensory and detoxification gene repertoires

One of our goals was to determine if chemosensory and detoxification gene gain, loss, and/or turnover (i.e., cumulative gains+losses) rates in *S. flava* deviate from those observed across non-herbivorous *Drosophila.* We chose seven *Drosophila* species for comparison with *S. flava* (Fig. S1a) which have high-quality genome assemblies, utilize a range of different feeding substrates with different degrees of specialization, and span the breadth of evolutionary distances from *S. flava* across the *Drosophila* phylogeny. We curated the complete set of genes within nine chemosensory and detoxification gene families (Cytochrome P450s, CYP450; Glutathione *S*-transferases, GSTs; UDP-glycosyltransferases, UGTs; Gustatory Receptors, GRs; Ionotropic Receptors, IRs; Odorant Binding Proteins, OBPs; Pickpocket Proteins, PPKs; Transient Receptor Potential channels, TRPs) in each genome, using stringent automated quality controls and manual inspections to minimize artifactual gene absences, duplications, and collapsed paralogs that could drive spurious patterns in *S. flava.*

Contrary to the expectation that herbivory spurs the expansion of gene families that interact with plant secondary compounds, *S. flava* has the smallest repertoire of detoxification genes, and nearly the smallest repertoire of chemosensory genes (with only one more gene than *D. virilis),* observed across the eight genomes (Fig. 3a, upper panel). A trend toward gene loss is recapitulated at the level of individual gene families: none of the curated gene families in *S. flava* exceed the range of gene counts observed in the other *Drosophila,* and most are near or below the lowest observed count (Fig. 3a, lower panel). This pattern is consistent with the neural limitation hypothesis, which posits that specialist herbivores have fewer decisions to make regarding where to lay eggs and feed compared to generalists, are more efficient when making such decisions, and may, accordingly evolve a simpler neural system. One prediction of this hypothesis is that specialists encode fewer chemosensory receptor types (37–39). This may extend to detoxification genes as well, given that fewer types of molecules are encountered by specialists owing to their relatively narrower diet breadth.

**Figure 3.**
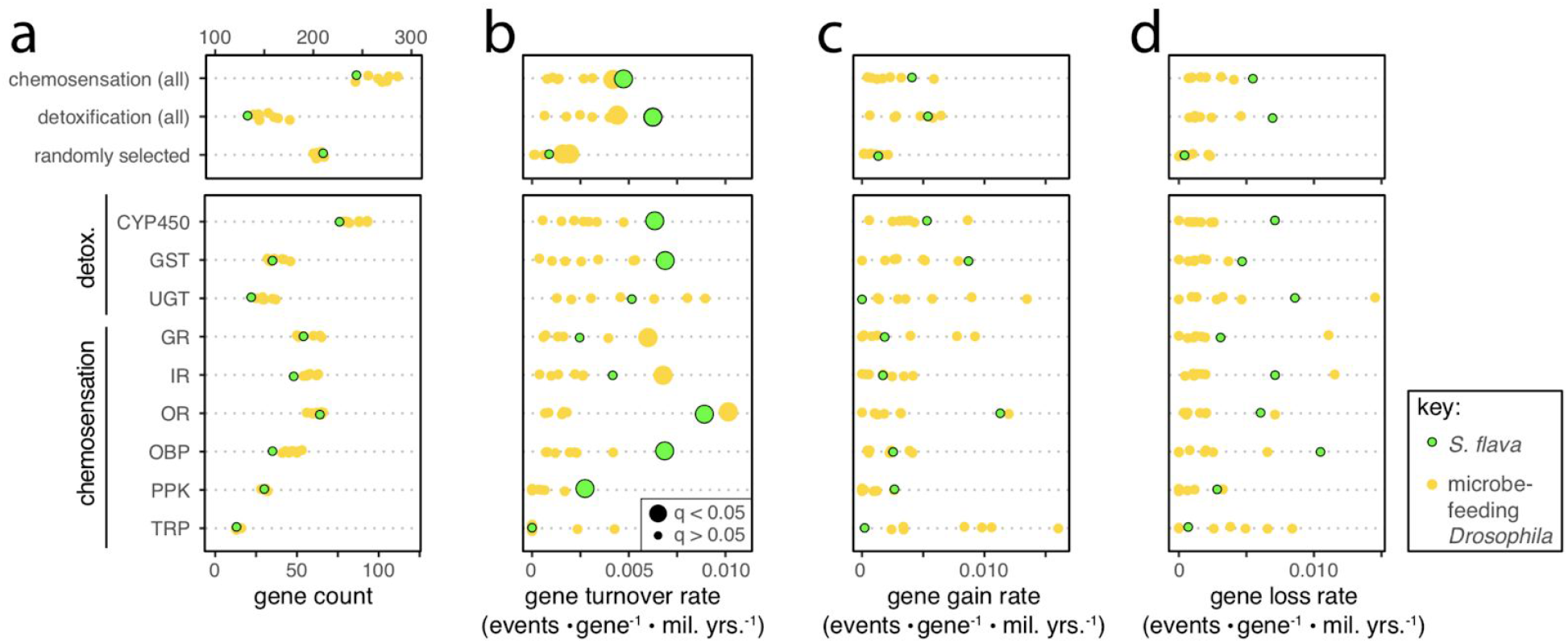
Elevated gene turnover rates within chemosensory and detoxification gene families in *S. flava*. All four panels share the same y-axis, the lower box in each panel shows results for nine annotated gene families, and the upper box shows results for genes grouped into three broad categories: chemosensory genes, detoxification genes, or genes randomly selected from the full genome (N=200 clusters of orthologous genes), **(a)** The number of genes per family in *S. flava* and seven non-herbivorous *Drosophila,* **(b)** The rate of gene turnover, a single parameter for the cumulative rate of gene gain and loss events, was estimated for each of the eight taxa using maximum likelihood. Turnover rate was further decomposed into separate rates of gene loss **(c)** and gain **(d).** In panel **(b),** lineage-specific rates that are significantly accelerated relative to the rest of the phylogeny were identified by comparing two models--a null model with a rate shared across the entire phylogeny, and an alternative model with a different rate in the focal lineage--through likelihood ratio tests at an analysis-wide false discovery rate of 5% (i.e., *q* < 0.05). Note that gene turnover rates were estimated in separate models than gene gain and loss rates, so the inferred gain and loss rates do not directly sum to the inferred turnover rates. Abbreviations: CYP450, Cytochrome P450; GST, Glutathione S-transferase; UGT, UDP-glycosyltransferase; GR/IR/OR, Gustatory, Ionotropic, and Olfactory receptors; OBP, Odorant binding protein; PPK, Pickpocket (DEG/ENaC); TRP, Transient receptor potential channel.

Comparisons of gene family sizes, however, offer only superficial insights into the evolutionary dynamics of these families across the phylogeny. Gene families could be functionally remodeled through accelerated gene gains and losses, even in the absence of overall changes in total gene number. We tested if the evolution of herbivory in *S. flava* was coupled with lineage-specific acceleration in the cumulative rate of gains and losses (i.e., gene turnover) using a maximum likelihood, phylogenetic approach that incorporates gene-level orthology relationships. *S. flava* exhibited the fastest turnover of both detoxification and chemosensory genes (Fig. 3b; analysis-wide false discovery rate = 5%).

Gene turnover can be further decomposed into rates of gain and loss. Most gene families undergoing accelerated turnover in *S. flava* exhibited atypically high rates of both gain and loss relative to other *Drosophila* (CYP450s, GSTs, ORs, and PPKs). On the other hand, OBPs exhibited only a strongly accelerated rate of gene loss.

A major implication of these findings is that studies that compare gene counts between herbivorous and non-herbivorous taxa may overlook important dynamics of gene family remodeling. When rates of gene gain and loss increase together, their net effect may be only a modest change in gene overall gene family size. In the case of *S. flava,* the accelerated gene turnover associated with herbivory only manifests as a small reduction in overall gene counts.

### Accelerated duplications and evolution of a key detoxification gene

To functionally test how plant-derived chemicals drove gene family dynamics in *S. flava,* we first focused on its interactions with the glucosinolate (GSL)-myrosinase system (the “mustard oil bomb”; (40)), which is the principal chemical defense system in the Brassicales. GSLs are amino-acid derived molecules that are hydrolyzed upon tissue damage, forming highly electrophilic and insecticidal isothiocyanates (ITCs or mustard oils) (41, 42). Ingestion of mustard oils or exposure to mustard oil vapor slows growth and is highly toxic to insects, including *D. melanogaster* (43, 44). Many insects specializing on Brassicales plants, however, have evolved physiological mechanisms to avoid toxic effects of mustard oil exposure during feeding, usually by preventing the hydrolysis of GSLs into mustard oils altogether and even sequestering intact GSLs (45–51). Although these cases reveal exquisite adaptations for disarming the “mustard oil bomb”, they do not necessarily reveal how host-switching events to GSL-bearing Brassicales plants actually unfold. While such adaptations may rapidly evolve, it is possible that the first stages of adaptation to GSL-bearing plants proceeds in a more gradual fashion, whereby natural selection leverages pre-existing ITC detoxification systems, and standing allelic variation or copy number variation therein, to begin an adaptive walk. Such is the case for generalists, from lepidopterans (52) to drosophilid flies, including *D. melanogaster,* which was first shown in radiolabeling experiments (53) to detoxify ITCs using the glutathione-dependent mercapturic acid pathway, the first step of which (dithiocarbamate formation) is catalyzed by GSTs.

Surprisingly, our previous metabolic profiling revealed that ITCs accumulate in *S. flava* tissue during feeding on mustards (53). Although physiological exposure to mustard oils is costly to *S. flava*—ingestion of GSLs slows development time (Fig. 4a), reduces weight gain, and induces physiological stress (23)—the detrimental impacts of GSLs on *S. flava* growth and performance are small compared to insects that do not specialize on GSL-bearing plants (e.g., (54)). Together, these patterns suggest that dietary GSLs likely presented a major barrier to the evolution of herbivory and mustard specialization in the ancestor of *S. flava,* and that an efficient mechanism of mustard oil detoxification likely played a key role in overcoming this barrier. Our metabolic profiling (53), after feeding *S. flava* radiolabeled glucoraphanin, revealed that all four Brassicales specialist *Scaptomyza* species tested metabolize mustard oils using the mercapturic acid pathway (Fig. 4b), a versatile pathway for detoxification of electrophilic molecules that is present across eukaryotes and predated the evolution of herbivory (55). GSTs catalyze the first step of the mercapturic acid pathway for diverse substrates, including mustard oils.

**Figure 4.**
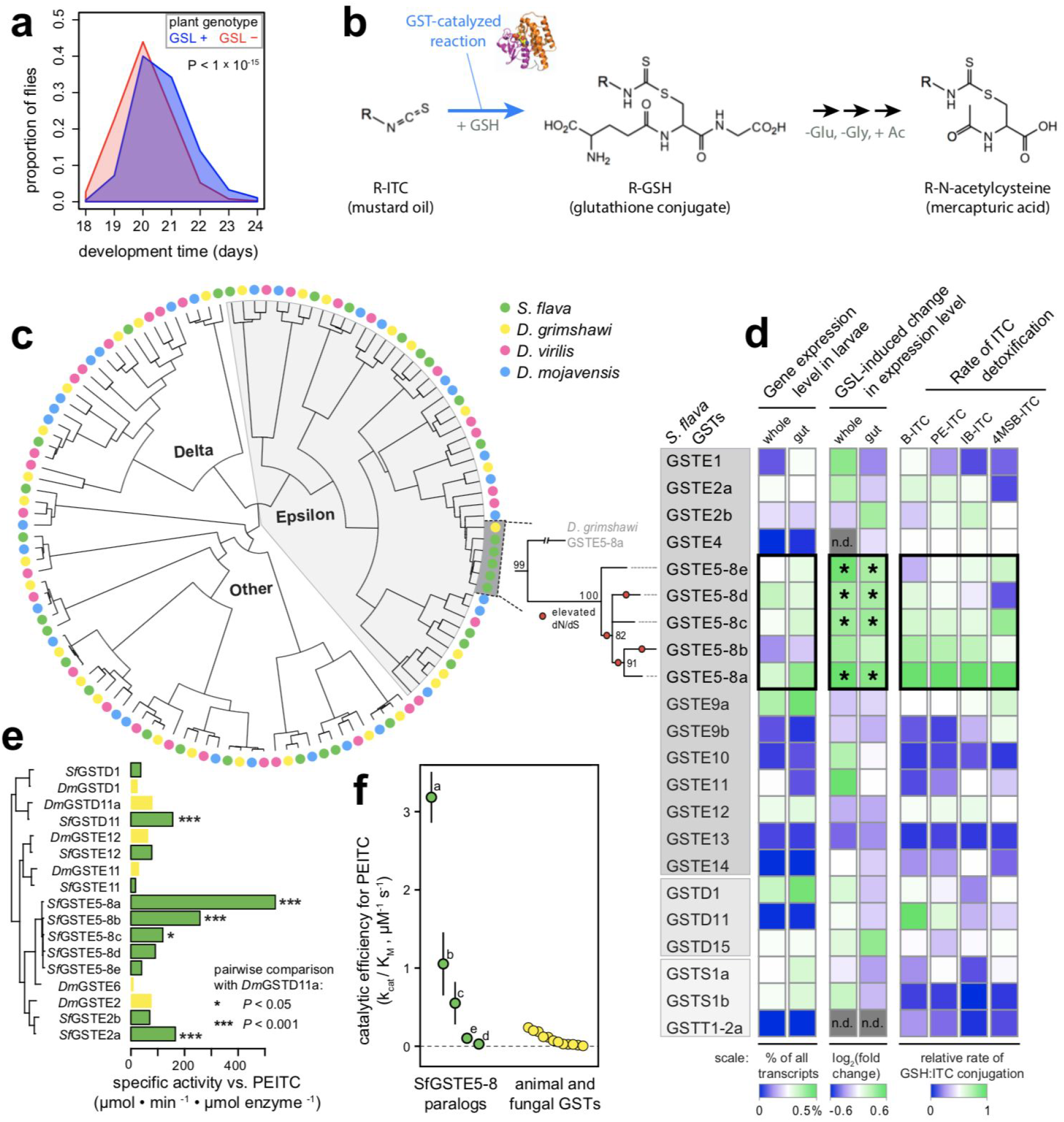
*S. flava* harbors a lineage-specific expansion of GSTs which are induced by, and efficiently detoxify, the primary defensive toxin from its host plants. **(a)** Egg-to-adult development time on *A. thaliana* plants is accelerated when the aliphatic and indolic GSL biosynthesis pathway is knocked out (GSL-mutant: *myb28 myb29 cyp79b2 cyp79b3)¦* N = four cages per plant genotype, 3016 total flies, **(b)** GST enzymes catalyze the conjugation of mustard oils to glutathione (GSH), the first step in detoxification through the mercapturic acid pathway in *S. flava.* Adapted from (53). **(c) Left:** Maximum likelihood amino acid phylogeny of cytosolic GSTs in the subgenus *Drosophila.* **Right:** Maximum likelihood nucleotide phylogeny of *S. flava GstE5-8* genes, the largest lineage-specific GST expansion in *Drosophila.* Bootstrap support is shown at each node, and branches with significantly accelerated d/V/dS are marked with circles, **(d) Left:** Expression level of *Gst* genes in larvae fed on *A. thaliana.* The percentage of all sequenced transcripts expressed from each gene was inferred from Illumina RNAseq and is directly proportional to RPKM. **Center:** Difference in expression level in larvae feeding on GSL+ vs. GSL-plants. Expression was measured through Illumina RNAseq in two separate experiments: whole larvae were reared from egg on either GSL+ or GSL-plants and profiled after 5 days (23), and larval guts were profiled from 7-day old larvae reared on GSL-plants, transferred to GSL+ or GSL-plants, and profiled after 12 hours. “*” indicates differentially expressed genes relative to larvae that fed on GSL-plants. **Right:** Specific activity against ITCs measured in vitro under physiologically relevant conditions for heterologously expressed and purified GSTs from *S. flava.* **(e)** Specific activity against PEITC. The enzymes chosen for this assay were among the five enzymes in each fly species with the highest specific activity against PEITC in either the initial GST-wide activity assays in *S. flava* or in a previous GST-wide activity assay in *D. melanogaster (57*). **(f)** *k_cat_/K_M_*, which approximates catalytic efficiency under physiological conditions, for *S. flava* GSTE5-8 enzymes +/− SEM (labels refer to paralogs a-e; this study) compared to published estimates for other taxa from the literature (Table S5). Abbreviations: B, benzyl; PE, phenylethyl; IB, isobutyl; 4MSB, 4-methylsulfinylbutyl.

We next investigated if the presence of mustard oils in the diet of *S. flava* was coupled with evolution of its GST gene repertoire. Through a candidate gene approach, we previously found (53) that rapid accumulation of amino acid substitutions following a gene duplication event caused a highly conserved GST (*GstD1*) to more efficiently detoxify mustard oils in *S. nigrita,* a close relative of *S. flava* (Fig. 1c). Variation in amino acid sequence among paralogous GST copies also leads to marked differences in the rate of mustard oil detoxification in humans (56), although it is unknown whether this variation is adaptive. We therefore expected rates of gene duplication and molecular evolution in genes encoding GSTs to be elevated in *S. flava* relative to non-herbivorous drosophilids.

Although the number of GST genes in the *S. flava* genome fell within the range observed in other *Drosophila,* two patterns suggested the evolution of herbivory has spurred remodeling of its GST repertoire. First, as noted earlier, GST gene turnover rates are significantly accelerated relative to *Drosophila* as a whole, driven by the highest lineage-specific rates observed for both gene gain and loss (Fig. S1). Second, nearly one-third of all GSTs in *S. flava* have accumulated amino acid-changing substitutions at a significantly faster rate (*dN/dS*) than their orthologs in other lineages (Fig. S2). This pattern was driven largely through rapid evolution of duplicated GSTs (median fold increase in *dN/dS* in *S. flava =* 1.15 for single copy GSTs and 2.33 for duplicated GSTs).

The largest lineage-specific expansion of a single GST observed within the subgenus *Drosophila* occurred in *S. flava,* coupled with the rapid evolution of many of the descendant paralogs (Fig. 4c). We refer to this expanded clade of GST genes as *GstE5-8* genes, based on homology to a clade of GSTs containing *GstE5, GstE6, GstE7,* and *GstE8* in *D. melanogaster* (Fig. S3a). *D. grimshawi,* the closest relatives of *S. flava* included in our analysis, each have two single *GstE5-8* copies, whereas *S. flava* has five. Gene tree-species tree reconciliation (Fig. S3b) and mutation rate-calibrated estimates of pairwise divergence between paralogs (Fig. S3c) indicated that all lineage-specific *GstE5-8* genes in *S. flava* arose from a single ancestral gene after splitting from the ancestor of its non-herbivorous sister lineage (*Scaptomyza pallida),* broadly concurrent with the timing of the evolution of herbivory and feeding on Brassicales plants ca.10 million years ago.

To identify specific GSTs at which rapid molecular evolution was likely driven by adaptation to dietary ITCs, we conducted a transcriptome-wide gene expression experiment and *in vitro* enzyme function study for all cytosolic GSTs (delta, epsilon, sigma, and theta class) in *S. flava.* First, we used transcriptome sequencing to compare GST gene expression in the presence or absence of GSLs for both whole larvae (23) and larval guts (this study), enabled by *Arabidopsis* mutant lines that allowed us to rear larvae in plants differing in the presence or absence of a functional GSL biosynthesis pathway. Only the expression of *GstE5-8* genes, which together comprise > 1% of all mRNA transcripts in the gut (Fig. 4d, left), was significantly induced by dietary GSLs (Fig. 4d, middle). Second, we profiled the *in vitro* rate of mustard oil detoxification using pure solutions of each *S. flava* GST, obtained by heterologous expression and sepharose purification. GSTE5-8 enzymes from *S. flava* had exceptionally high activity against mustard oils (Fig. 4d, right). Three GSTE5-8 enzymes, as well as two other *S. flava* GSTs, detoxified mustard oils more efficiently than the most efficient GST identified in a GST-wide screen in *D. melanogaster* (57) (Fig. 4e). The most active enzyme, SflaGSTE5-8a, has a catalytic efficiency for dithiocarbamate formation that is an order of magnitude higher than any previously characterized GST from metazoans (Fig. 4f).

Together, these results reveal that constant exposure of *Scaptomyza* to mustard oils, a novel dietary toxin for drosophilids, was coupled with rapid rates of sequence evolution and gene duplication in the GST gene family. This process gave rise to a bloom of highly expressed detoxification enzymes with exceptional affinity for mustard oils—thus providing a direct link between the evolution of herbivory and the putatively adaptive remodeling within a detoxification gene family in *S. flava.*

### Gene blooms, not gene family expansions, as a consequence of herbivory

Given that the largest bloom of GSTs across *Drosophila* occurred in *S. flava* and gave rise to highly efficient enzymes to detoxify its major host plant-derived toxin, we reasoned that functionally-important gene blooms may extend to other gene families. To search for the most extreme cases, we binned genes into orthologous groups that likely each originated from a single copy in the ancestor of all *Drosophila,* and determined if *S. flava* had the largest expansion of any single orthologous gene group within any other chemosensory or detoxification gene families.

We found that *S. flava* also exhibits the largest bloom of any CYP450, a lineage-specific increase increase to six *Cyp6g1* genes (equaled only by the six *Cyp4p* genes in *D. grimshawi*). This is notable because *Cyp6g1* has repeatedly undergone lineage-specific expansions in other *Drosophila* (to as many as three gene copies) (58), coupled with functional divergence that includes the acquisition of resistance to different insecticides (28, 29, 59). Intriguingly, CYP450s in spider mites likely confer experimentally evolved resistance to indolic GSLs (60), a subset of GSLs that are widespread among the Brassicales and whose insecticidal activity does not arise from conversion to mustard oils (60). Thus, while the substrates for *Cyp6g1* genes in *S. flava* are unknown, GSLs may play an important role in this gene bloom as well. Regardless of the substrates driving this pattern, however, the bloom of *Cyp6g1* in *S. flava* lends further support to the hypothesis that the evolution of herbivory does not necessarily require dramatic changes to detoxification repertoires, but instead can rely on expansions of the same functionally versatile genes that allow their non-herbivorous relatives to cope with xenobiotic challenges.

*S. flava* also had the largest observed bloom of any OR across the drosophilids in our comparison, an increase to seven *Or98a* genes (matched only by *Or42b* in *D. grimshawi*). Like *Cyp6g1, Or98a* likely plays a key role in chemical niche adaptation in the non-herbivorous relatives of *S. flava:* it is expressed in one of the few neurons with altered odorant sensitivities among populations of *D. mojavensis* that specialize on rots of different cactus species (61). Given that *Or98a* is required for mating behavior in *Drosophila* (62), this bloom may be important for the evolution of a novel mating niche associated with living plants in *S. flava.* This adds to the evidence found previously, and verified by the present genome assembly, that ORs play a key role in behavioral shifts in the evolution of herbivory: *S. flava* also gained two additional copies of the broadly-tuned *Or67b* (Fig. S4) that exhibit signatures of positive selection (32), and lost three canonical ORs (*Or22a, Or42b,* and *Or85d*) associated with perception of yeast volatiles (32). These patterns are consistent with those in other specialized insects and reveal that chemoreceptor losses and blooms may be predictable when niche shifts occur (38, 63).

How do these gene blooms inform our understanding of the effects of diet on adaptive gene family evolution? Previous work suggested the evolution of one of the largest detoxification gene families across insects, the CYP450s, may generally be driven by stochastic birth-death processes, and not through chemical co-evolutionary (*sensu lato*) interactions between insects and plants (64). However, new studies suggest that this finding obscures potentially adaptive “blooms” that are more nuanced, localized phylogenetically and driven by the particular niches of the insects involved. Specifically, studies on insects with well-defined interactions with plants, including shifts from carnivory to folivory in bees and variation in diet breadth across *Drosophila* and Lepidoptera species, largely refute a purely stochastic birth-death hypothesis (19, 58, 63–65). We found a similar pattern of fewer CYP450 gene numbers in *S. flava,* likely due to a narrowing of their dietary niche, coupled with high turnover rates (high gain + loss rate) and a modest, but biologically significant bloom in the canonical xenobiotic resistance gene *Cyp6g1.* A similar pattern is observed with respect to GSTs: although there was no large shift in overall gene number, a bloom of genes also evolved in the *GstE* subfamily that we functionally characterized to be differentially regulated by dietary ITCs and were the most efficient in detoxifying mustard oils of any GST known from any animal. It is unlikely that these blooms were predominantly stochastic.

The targeted gene expansions observed in chemosensory and Phase I and II xenobiotic metabolism ontologies are also consistent with the long-standing prediction that chemical co-evolutionary arms races should drive reciprocal change in host plants and the specialized insects that attack them (10, 14). Diffuse co-evolution can clearly promote gene duplication and subsequent neofunctionalization or even concerted neofunctionalization within detoxification gene families in mustard specialists (13, 49, 50, 53).

### Drivers of gene loss

The evolution of herbivory is not just a shift to a new niche, but also necessarily a transition away from an ancestral one. While gene losses could arise through relaxed selection on ancestrally important traits, they could also confer adaptations for plant feeding. Although disentangling these two mechanisms to explain accelerated gene loss in *S. flava* is not possible within the current study, the striking loss of OBPs, which outpaced gene losses in any other chemosensory gene family, offers an intriguing avenue for future investigation. The majority of OBP losses in *S. flava* (seven of nine) were within the Plus-C OBP subfamily that is otherwise highly conserved within *Drosophila*? for comparison, among the three taxa we examined that were most closely related to *S. flava,* there are only two combined losses of Plus-C OBPs. Plus-C OBPs encode six additional cysteine residues compared to other OBPs (105). The additional cysteines may render them more vulnerable than typical OBPs to damage by mustard oils, which are known to attack disulfide bridges between cysteines (106) and to which OBPs secreted into the sensory lymph of sensilla would be directly exposed. This raises the possibility, although speculative, that bursts of gene losses, like gene blooms, may be driven in herbivores by functional properties shared among genes.

### Robustness of findings to biological and bioinformatic confounding

In comparative genomics studies, misleading conclusions can arise when the variable of interest (e.g., the evolution of herbivory) is biologically confounded with a causal variable (e.g., demographic events) (20), or with a bioinformatic approach that lacks sensitivity or imparts lineage-specific biases (66). We considered three strong sources of potential confounding in *S. flava*—shifts in demography (e.g., population size); artifactually missing, collapsed, or duplicated regions of the genome assembly; and incomplete gene annotations. For reasons outlined below, we conclude they are unlikely to explain exceptional patterns of detoxification and chemosensory gene evolution.

Demographic processes, such as population bottlenecks, can weaken the efficacy of natural selection. This could lead to an accelerated fixation of slightly deleterious gene gains and losses. Three lines of evidence from our analyses suggest demographic processes are unlikely to explain the results. First, we curated a randomly selected set of orthologous gene clusters to infer genome-wide rates of gene turnover. Both gene and loss rates in *S. flava* fell squarely within the range of rates observed in other *Drosophila* (Fig. 3). Second, we inferred the level of nucleotide diversity (π) in an *S. flava* population using pooled whole genome sequencing. Autosomal nucleotide diversity, which is proportional to effective population size, was similar between a single population of *S. flava* (π = 0.0056) and *D. melanogaster* (the DGRP; π = 0.0060 (67)). The relationship between physical distance and linkage disequilibrium was also similar between the two species and decayed quickly (Fig. S5), consistent with sharing similarly large effective population sizes (107). Third, we estimated the proportion of the genome composed of repetitive elements, which is thought to be sensitive to demographic shifts (68). Repeat content in *S. flava* is within the range observed in other Drosophila (Table S6).

Given that the *S. flava* genome was sequenced, assembled, and annotated with a different approach than the *Drosophila* genomes included in our study, we used a number of different strategies to minimize artifactual gene gains and losses. First, we mapped unassembled short reads back to the genome assembly and quantified read depth across the curated genes. Cumulative read depth was highly predictive of the final number of paralogous copies annotated for each gene (Fig. S6). Second, for all genes with lineage-specific gains and losses in the original *S. flava* assembly, we performed manual re-curation in a second, independent, long-read genome assembly. The few discrepancies between assemblies were all clearly resolved as cases of artificially collapsed or duplicated scaffolds in one of the assemblies on the basis of abnormally high or low read depth. Third, we identified the genomic regions in both assemblies syntenic to the likely ancestral locations of genes lost in *S. flava.* We found contigs spanning across 95% of these regions, yet failed to find any functionally-intact genes missed by our automated and manual curations. Together with the high completeness metric for core dipteran genes (Table S3-S4) and the lack of abnormally high rates of gene duplication or loss in the set of randomly curated genes, we conclude that assembly and annotation biases are unlikely to explain accelerated gene family dynamics in *S. flava.* This extends to gene blooms in *S. flava,* which were each unequivocally supported by agreement between the two genome assemblies (Fig. S4), read depth analyses, and (for *GstE5-8* paralogs) targeted amplification and Sanger sequencing of cDNA.

### Conclusions

Genomic comparisons among anciently evolved herbivores and distantly related non-herbivores, or across herbivorous lineages, have uncovered striking expansions and losses of genes involved in chemosensation and detoxification in arthropods (e.g., (19, 32, 65, 69, 70). Yet the lack of dense sampling among closely related taxa has, in many cases, precluded pinpointing the timing of these changes relative to the evolution of herbivory. Here, the depth of genomic resources from *Drosophila* illuminated accelerated gain, loss, turnover, and blooms of detoxification and chemosensory genes in one of their herbivorous relatives. By linking these dynamics to the transition to herbivory in a phylogenetic context, the genome of *S. flava* lends powerful support to the hypothesis that the chemical composition of plant tissues drives herbivore genome evolution, an idea at the heart of early theories on species interactions that motivated the development of co-evolutionary theory (10, 71–73). Similar comparative approaches in other lineages promise to reveal the extent to which the genomic changes tied to herbivory in *S. flava* reflect general strategies underpinning the evolution of herbivory-and the bursts of biodiversity that follow.

## METHODS

### Herbivory experiments and isotope profiling

#### Oviposition preference assays

*S. flava* has exclusively been reared from living plants when collected in nature (23, 26, 53). To determine if *S. flava* occupies a broader feeding niche than reported previously, we offered adult female flies a choice between living plants, decaying plants, and yeast-seeded media. Six week old *A. thaliana* (Col-0) were provided as living or rotting plants. To facilitate decay, plants were frozen overnight and subsequently placed in a warm, moist environment for three days before use. Standard *Drosophila* media seeded with *Saccharomyces cerevisiae* (Fleischmann’s) was prepared in petri dishes. The three food substrates were placed randomly in a 30 x 30 cm mesh cage. 6-8 day old flies from an outbred colony collected near Dover, NH, USA were placed inside the cage, and eggs were counted from each substrate after 24 hours. The experiment was conducted at 22°C and 60% humidity.

#### Viability assays

To determine if *S. flava* requires living plants to complete development, larval viability was assayed on living plants, decaying plants, and yeast-seeded *Drosophila* media, prepared and maintained as described above. Because *S. flava* will only oviposit on living plants, 3-5 day old larvae were transferred from *A. thaliana* (Col-0) to each substrate. Nine replicates of six larvae were set up for each substrate, and emerged adults were censused and removed at 13, 20, and 27 days.

#### Isotope profiling

*S. flava* was reared in replicate cages with ample Col-0 rosettes. Plant tissue was collected immediately prior to exposure to adult *S. flava,* and again after 10 days of feeding using leaf tissue adjacent to any mines. 25 third instar larvae were collected for each of three replicates at 10 days and at 21 days. To compare to the microbe-feeding *Drosophila,* wild *Drosophila melanogaster were* allowed to colonize three diets differing in the photosynthetic (i.e., C-isotope discrimination) ability: *Cucumis melo,* melon (C3 pathway), *Opuntia* sp., fruit (CAM), and *Zea mays,* maize (C4 pathway). Substrates were placed outside in Tucson, AZ, USA to allow colonization by flies and were subsequently inoculated by fungus. Diet, third instar larvae, and adults were collected from each substrate. Three replicates for all species-diet combinations were used in the analysis. Immediately after collection, tissues were dried at 50°C, ground to a fine powder, and combusted using an Elemental Combustion System (Costech Analytical Technologies, Valencia, CA) coupled to a continuous-flow gas-ratio mass spectrometer (Thermo Finnigan Delta PlusXL, San Diego, CA). δ^13^C, δ^15^N, C, and N were quantified using acetanilide standard for elemental concentration, NBS-22 and USGS-24 for δ^13^C, and IAEA-N-1 and IAEA-N-2 for δ^15^N. Precision is better than ± 0.10 for δ^13^C and ± 0.2 for δ^15^N (1 s.d.), based on repeated internal standards.

### Species diversity of herbivorous *Scaptomyza*

*Scaptomyza* feeding on Brassicales spp. were collected in CA, USA, in 2017-2018. DNA was extracted and *COI* was sequenced as described previously (25, 53). Additional *COI* sequences were retrieved by the following accession numbers: *S. graminum* (LC061494.1); *S. nigrita* (USA) (KJ943846.1); *S. nr. nigrita* (OR) (KJ943848.1); *S. nr. nigrita* (NV) (KJ943847.1); *S. nr. flava* (AZ) (KJ943850.1); *S. flava* (USA) (JX160022.1); *S. flava* (NL) (KJ943851.1); *D. grímshawi* (BK006341.1:1455-2990); *S. apicata* (JX160024.1); *S. hsui* (KC609729.1). *COI* sequences were aligned using MUSCLE (74) and visually inspected, and 765 bp were retained for further analysis. A maximum likelihood phylogeny was constructed using the Tamura-Nei model (75) with 500 bootstraps in MEGA v10.0.4 (76). Initial tree(s) for the heuristic search were obtained automatically by applying Neighbor-Join and BioNJ algorithms to a matrix of pairwise distances estimated using the Maximum Composite Likelihood (MCL) approach. The topology with highest likelihood score was retained and rooted at the bifurcation between Caryophyllaceae specialist *S. graminum* and the rest of Brassicales-feeding *Scaptomyza* following (32).

### Genome sequencing, assembly, and annotation

#### Biological material and sequencing

The sequenced line (OHI-9) of *S. flava,* derived from a laboratory population initially collected in Belmont, MA, USA in 2008, was passaged through 10 generations of single pair mating on *A. thaliana* Col-0 plants. Paired-end 180 bp and 200 bp insert libraries and 3 kbp and 5 kbp mate pair libraries from OHI-9 female flies were sequenced with 100 bp read length on an lllumina Hiseq 2000 at the University of Arizona. Reads were quality filtered and Illumina TruSeq3 adapters were removed using Trimmomatic vθ.35 (77) with the following parameters: “LEADING:10 TRAILING:10 SLIDINGWINDOW:4:15 MINLEN:99”.

#### Assembly and annotation

Read pairs that survived quality filtering were subsampled to an estimated ~90x coverage of the S. flava genome and assembled using ALLPATHS-LG (78) on the XSEDE high performance computing system. Contigs were extended and ambiguous regions were resolved iteratively using GapCloser (79) to produce the primary assembly used for all analyses (unless otherwise noted). Prior to annotation, repeat regions were masked using RepeatMasker (80) with the *Drosophila* repeat library. Protein-coding genes were annotated using MAKER2 (81), with the *S. flava* transcriptome (23) and predicted gene sequences from 12 *Drosophila* species (FlyBase release 2013_06) provided to inform gene models. We recovered 17,997 genes in our liberal gene set (annotations predicted by Augustus) and 12,993 genes in our high confidence gene set (annotations supported by multiple lines of evidence in MAKER2). The proportion of core dipteran genes recovered in our assembly was determined with BUSCO v3.0.2 (82).

We subsequently used SSPACE-Longread (83) to perform further scaffolding using PacBio reads. Because this did not improve BUSCO benchmarking statistics for genome completeness or fragmentation, we did not use this scaffolded assembly for any analyses, but will make it available for applications that benefit from longer scaffold lengths (e.g., GWAS).

#### Repetitive element content

Repeat identification was carried out using both homology-based and ab-initio approaches. We used the Drosophila RepBase repeat database for the homology-based annotation (http://www.girinst.org/repbase; update 20150807) within RepeatMasker version open-4.0.6 (80). The RepeatMasker option-gccalc was used to infer GC content for each contig separately to improve the repeat annotation. Ab-initio repeat finding was carried out using RepeatModeler version 1.73 (http://repeatmasker.org/RepeatModeler.html).

#### Generation of a long read genome assembly

To assist in validation of gene models, an independent long read assembly was generated from a partially inbred *S. flava* colony. Please see the Supplementary Note for details.

### Gene family evolution

#### Initial model curation

An iterative curation strategy was used to identify the complement of chemosensory and detoxification genes in each genome. First, all protein sequences in each family from *D. melanogaster were* queried against protein annotations for each member of the subgenus *Drosophila* using BLASTP (84). Putative gene family members (e-value < 1e-3) were retained. The resulting collection of genes was then queried against the annotated *S. flava* protein models, retaining putative *S. flava* orthologs (e-value < 1e-3). Amino acid sequences were aligned using MUSCLE (74) and manually inspected.

To retain only true gene family members, a maximum likelihood phylogeny was constructed from these sequences using RAxML blackbox (85) with default settings. Identical sequences from separate scaffolds were flagged as putative assembly errors and reduced to a single copy. Alignments of gene models of *S. flava* genes significantly deviating in length from their *D. melanogaster* orthologs were manually inspected, and *S. flava* gene models were corrected when necessary using homology to the annotated genes in other species. Finally, genes from all species were realigned and a maximum likelihood tree was constructed, using the same methods described above.

#### Validation of gene curations

We manually re-inspected all cases of lineage-specific gain and loss (hereafter “genes of interest”, GOIs) in *S. flava* using a number of additional approaches.

To validate lost genes GOIs, we performed an additional TBLASTN search for each GOI in *S. flava,* as well as genes assumed to be proximal to GOIs (PGOIs) based on their location in *D. grimshawi* or *D. virilis.* These searches used the predicted orthologs in *D. grimshawi* (as determined using the phylogenies described above) as queries. We performed this search in the original *S. flava* ALLPATHS assembly as well as a long-read assembly (see results) that has not yet been released but is available upon request. Genes of interest were considered truly absent if TBLASTN searches of both *S. flava* genome assemblies as well as a de novo *S. flava* transcriptome assembly (23) yielded no hits. In some cases, homology between conserved protein domains resulted in weakly supported hits to the *S. flava* genome and/or transcriptome, in which case the aligned region was extracted, translated, and BLASTed against the NCBI nr database.

The gene was considered lost if the output showed stronger homology to genes other than the GOIs. To avoid confirmation bias, we also did this with the *S. flava* PGOIs to ensure their identity matched the expected ortholog in *D. grimshawi.* The absence of the GOIs, coupled with the presence of 95% of the PGOIs, in two independent *S. flava* assemblies (as well as the transcriptome) strongly support that the GOIs are truly lost and are not an artifact of missing scaffolds in the ALLPATHS genome assembly. We implemented a similar BLAST-based search of both genomes to validate genes that underwent lineage-specific expansions in *S. flava.*

As further validation, we inspected read depth for each curated gene. A subset of the paired-end Illumina reads (200 bp insert library, ~50x coverage depth) were mapped back to the ALLPATHS assembly using BWA mem (86), and read depth was calculated using SAMtools depth (87). The depth metric for each gene model was assigned as its median read depth across all sites, and genes for which this depth metric deviated by more than 15x coverage from the median across all curated genes (a deviation of ~30%) were flagged. These gene models were then manually inspected, following the same approach as genes with lineage-specific gains or losses.

Finally, to guard against errors in our curation of the published *Drosophila* genomes, we compared our gene curations to those from published studies (58, 88, 89) and those inferred in OrthoDB (90). For any discrepancies, we performed comprehensive TBLASTN searches against the relevant genome assemblies to search for the full complement of orthologous genes, re-curated gene models if necessary, and manually inspected aligned gene models. In a few cases, we discarded genes that had high similarity (>99% identity) and perfect synteny to another scaffold in the assembly, as has been standard in other studies in *Drosophila,* because these are likely artifactual duplicates. We also imported gene curations from the subgenus *Sophophora* (*D. erecta, D. ananassae,* and *D. pseudoobscura*) from these published orthology annotations directly, which we deemed reliable upon inspection because their annotation is facilitated by their close relationship to *D. melanogaster.*

#### Estimation of gene gain, loss, and turnover rates

Orthologous gene groups were designated from the curated gene set as monophyletic clades if they had > 70 bootstrap support and their phylogenetic topology suggested they were present as a single gene copy in the ancestor of the genus *Drosophila.* For poorly supported clades, orthology groups were assigned on the basis of previous orthology assignments (58, 88, 89) and syntenic relationships when bootstrap support was < 70. If an orthology group lacked members in either subgenus, and thus may be too narrowly defined, it was merged with the phylogenetically closest orthology group.

We also generated a set of 200 randomly chosen orthology groups to enable comparison between the curated gene set and a genome-wide background. Orthology relationships among protein-coding genes in *S. flava* and other *Drosophila* genomes (FlyBase release FB2013_06) were assigned by similarity clustering using orthoMCL v2.0.9 (91) with default parameters and an inflation value of 1.5. using the liberal *S. flava* gene annotation set. Genes gained or lost specifically in *S. flava* were validated in both assemblies using the TBLASTN-based approach described above.

Maximum likelihood rates of gene family gain, loss, and turnover were estimated for each gene family through independent runs of CAFE v4.0.2 (92). Global models with either a single turnover rate (λ-only) or a separate gain and death rate (λμ) were estimated, as were a series of models that allowed these parameters to differ between a single focal branch vs. the rest of the phylogeny (i.e., a branch model with separate foreground and background rates). Significance of each λ-only branch model was determined using a likelihood ratio test, with the global λ-only model serving as the null model. Significance of the λ-only branch models for GSTs were further validated by comparing the observed foreground λ for a given branch with the distribution of λ values estimated for that same branch in 2,000 pseudo-datasets, which were simulated under the best-fitting λ value from the global λ model for the corresponding gene family (e.g., under the assumption of no branch-specific deviations in λ). All models on simulated and real datasets were run in triplicate, and the iteration of each model with the highest likelihood was retained.

#### Molecular evolutionary analysis

For each group of orthologous GSTs, amino acid sequences from five taxa (*S. flava, D. grimshawi, D. mojavensis, D. virilis, and D. melanogaster*) were aligned using MUSCLE, manually inspected and corrected, and back-translated to CDS DNA alignments using RevTrans. Branch tests for accelerated *dN/dS* (93) were implemented for each terminal branch using the codeml module of PAML v4.5 (94) as in (53).

### Effects of glucosinolates on larval development and gene expression

#### Larval assays and tissue collection

To test the effect of GSLs on *S. flava* performance, we compared egg to adult development time on plants with and without GSLs. Adult females were allowed to oviposit on two mutant *A. thaliana* genotypes: *pad3* or *myb28 myb29 cyp79b2 cyp79b3 (95),* hereafter *“GKO”* for “GSL knockout”). *pad3* rather than wildtype plants were used as a control because they have indole and aliphatic GSL pathways intact but lack camalexins, a non-GSL defensive compound also pleiotropically absent in *GKO* mutants (96, 97). Cages contained ~40 plants each of a single genotype (N = 4 cages per genotype). After 8 hours, adult flies were removed and offspring were allowed to develop at 22°C under a 16:8 hour light:dark cycle. Cage positions on shelves were rotated daily to minimize positional effects. After two weeks, cages were monitored daily for emergence of adult flies, which were removed and counted.

In a separate experiment to detect GSL-induced changes in gene expression, adults were allowed to oviposit on *GKO* plants. After one week, larvae were transferred to feed for 12 hours on either *pad3* or *GKO* plants (N = 50 larvae per genotype). Larvae were initially reared on *pad3* because GSLs have strong effects on early larval development rate (23), so larvae reared from eggs on plants with and without GSLs quickly become out of sync developmentally, and could therefore exhibit developmentally-based changes in gene expression. 12 hours after transfer to the new host plant, larval guts were excised under a Stemi 2000-C microscope (Zeiss) and briefly stored at 4°C in RNAIater (Thermo-Fisher Scientific). RNA was extracted from the *pad3-* and GKO-transferred larval gut pools using a PureLink RNA Mini Kit (Thermo-Fisher Scientific).

#### RNA-sequencing and differential expression analysis

The GKO- and *pad3*-transferred larval gut pools were each sequenced to yield 100 bp paired-end reads on one quarter of an Illumina HiSeq 2500 lane at the University of Arizona. Over 50 million read pairs were mapped to *S. flava* gene models using TopHat2 v2.0.1 (98) with default parameters with the following exceptions: the maximum intron length was set to 50000 with a mate-pair inner distance of 100. We also used the same protocol to re-map single-end reads from a similar experiment using whole *S. flava* larvae and wild-type Col-0 (rather than *pad3) A. thaliana* (23). The number of reads mapping to specific loci were counted using HTseq (108).

Differentially expressed *Gst* genes were detected under a pairwise model using default settings in DEseq2 (99) to compare *pad3-*fed vs. GKO-fed larval guts and wildtype-fed vs. GKO-fed whole larvae. To increase power to detect differential expression for genes that show consistent expression trends between guts and whole larvae, P-values were merged across experiments using Stouffer’s Z method (100) with the R package metaP.

### GST activity assays

#### Heterologous expression and purification

Cytosolic GSTs from *S. flava* and *D. melanogaster were* PCR-amplified using cDNA generated from 7 day old larvae. *S. flava* were reared on *A. thaliana* (Col-0). Specific PCR primers were based at the 5’ and 3’ ends of the coding sequence. PCR products were first cloned into the TA cloning vector pGEM-T and sequenced. Sequence-validated products were then PCR amplified from the plasmid and cloned into different combinations of the BamHI, EcoRI, Xhol, or Sacl restriction sites in the plasmid pET28a (Novagen), depending on which restriction sites were present in each GST. This produced GSTs with an N-terminal 6His tag that was used for protein purification. GSTs were Sanger sequenced again, and validated GST-containing plasmids were transformed into *E. coli* BL21 chemically competent cells for protein expression.

Terrific broth (0.05-0.5 L) supplemented with 50 μg/mL kanamycin was used to culture *E. coli* BL21 cells containing each of the cloned GSTs. Cultures were grown at 37 C and shaken at 300 rpm until they reached an OD_600_ of 0.6-0.9 before induction of GST expression with 1 mM IPTG. The temperature was subsequently reduced to 30 C for 4-8 hours to facilitate protein expression. Cells were then pelleted and resuspended in 25-100 mL lysis buffer, pH 7.2-9.3 depending on the GST’s isoelectric point (50 mM NaPO4, 300 mM NaCl, 10% glycerol, 500 μg/mL lysozyme, 12 mM imidazole, 1x Protease Inhibitor Cocktail [Roche]). All subsequent steps were performed at 4 C. Cell lysate was disrupted by pulse sonication, and debris was removed by centrifugation. Cleared supernatant, containing soluble protein, was incubated with 500 μL of Ni sepharose resin (GE Healthcare Life Sciences) for 60 minutes and subsequently centrifuged to separate the sepharose resin from unbound soluble protein. The resin was washed 3x with wash buffer, pH 7.2-9.3 (50 mM NaPO4, 300 mM NaCl, 25 mM imidazole) to remove nonspecifically bound protein. GST proteins were then eluted with 10 mL of elution buffer, pH 7.2-9.3 (50 mM NaPO4, 300 mM NaCl, 250 mM imidazole). Eluted protein was buffer-exchanged to remove imidazole and concentrated using Amicon Ultra centrifugal filtration units with a 10 kDa molecular weight cutoff (EMD Millipore) and concentration buffer, pH 7.2-9.3 (50 mM NaPO4, 200 mM NaCl, 3 mM KCl, 10 mM KH2PO4). Purified protein concentration was determined using Bradford Reagent (Sigma Aldrich) with BSA as a standard. After determining protein concentration, glycerol was added to 50% final concentration and protein aliquots were stored at −80°C until use in specific activity and kinetic assays. Protein purity and expression was verified by SDS-PAGE electrophoresis. It was not possible to purify *S. flava* GSTT1/2b and *D. melanogaster* GSTD11b using these methods, but all other cloned GSTs were successfully purified and used in subsequent enzyme activity assays.

#### Enzyme activity assays

Assays were performed according to our previously described methods (53) with the following modifications: *[1]* The amounts of GST proteins to be used in each specific activity assay were determined empirically by testing several concentrations of each GST against each substrate to achieve a near-linear reaction rate, and varied from 0.33 μg to 20 μg GST per 300 μL reaction depending on the enzyme activity observed. *[2]* A saturating amount of glutathione (GSH) is required for kinetics experiments in which ITC is the rate-limiting substrate. This amount was determined through kinetics experiments using a fixed concentration of 2.5 mM CDNB and varying the GSH concentration from 25 μM to 20 mM. Among the GSTs tested, the *K_M_* for GSH ranged from 1.7 mM to 11.7 mM. Thus, 25 mM glutathione was used in all kinetics experiments.

### Population genomics

#### Pooled genome sequencing

A total of 45 *S. flava* larvae were collected from *Turritis* (formerly *Arabis) glabra* from a large field in Belmont, MA, USA in 2013. Each larva was collected from a separate plant individual to minimize relatedness. DNA was extracted from the pool of larvae using a DNeasy Blood and Tissue Kit (Qiagen). 100 bp paired-end sequencing was conducted on half of a lane on an Illumina HiSeq 2000 at the University of Arizona.

#### Read mapping

Reads were trimmed of adapters, and trimmed and filtered for quality with Trimmomatic v. 0.32 (77) using the following settings: ILLUMINACLIP:2:30:10, TRAILING:3, HEADCROP:2, SLIDINGWINDOW:6:15, and MINLEN:50. Retained reads from each population were then mapped to the *S. flava* reference genome first with BWA v. 0.6.1 (86) using the MEM algorithm, and then using Stampy v. 1.0.23 (101) using the-bamkeepgood reads option. The substitution rates supplied to Stampy were taken from previous Stampy runs and insert size statistics were estimated from previous mappings using Picard v. 1.107 CollectInsertSizeMetrics. Resulting SAM files were converted to BAM files using SAMtools v. 0.1.18 (87). BAM files were cleaned and sorted using Picard CleanSam and SortSam, and duplicates reads were marked and removed using Picard MarkDuplicates. Realignment around indels was performed using GATKv. 2.8-1 (102) RealignerTargetCreator and IndelRealigner. SAMtools was then used to remove unmapped reads, keep only properly mapped read pairs, filter for a mapping quality of 20, and create pileup files from each of the final BAM mapping files. Popoolation v. 1.2.2 (103) filterpileup-by-gtf.pl was used to filter the pileup files for the repetitive regions, identified as described previously. Reads were mapped to a mean coverage depth of 31,4x across the *S. flava* genome.

#### Genetic variation

Nucleotide diversity (π) was calculated per scaffold for four-fold degenerate sites using Popoolation Variance-sliding.pl for repeat-masked autosomal scaffolds (N = 819) greater than 20 kb in length. Parameters were: a ploidy level of 90, minimum allele count of two, minimum quality score of 20, minimum coverage of four, and a maximum coverage of 100.

#### Linkage Disequilibrium

LD was estimated from the 15 largest autosomal scaffolds. After filtering repeats from the BAM file using BEDTools v. 2.17.0 (109) intersect, SNPs were called using GATKv. 2.8-1 (102) UnifiedGenotyper with heterozygosity of 0.014, ploidy level of 90, two maximum alternative alleles, and a maximum coverage of 200. Preliminary SNPs were then hard filtered using GATKv. 2.8-1 VariantFiltration and SelectVariants. LDx (110) was used to obtain a maximum likelihood estimate of linkage disequilibrium (r^2^) with the following parameters: a minimum read depth of 10, a maximum read depth of 150, an insert size of 417, a minimum quality score of 20, a minimum minor allele frequency of 0.1, and a minimum read intersection depth of 11.

### Data availability

The primary assembly used for this analysis is available as GenBank assembly accession GCA_003952975.1. RNA-seq reads from midgut expression profiling are available as SRA accession SRX4512133. All other data and scripts will be made freely available upon acceptance of the manuscript in its final form, and are available from the corresponding authors upon request.

## ACKNOWLEDGMENTS

Financial support was provided to A.D.G. by the National Science Foundation (DEB 1405966 and a Graduate Research Fellowship); to A.D.G., B.G.H., and T.K.O. by the NSF IGERT program (traineeship DGE-0654435); to A.C.N.D., R.T.L., and D.H.H. by the National Institutes of Health (PERT fellowship 5K12GM000708-13); to P.D.N. by the United States Department of Agriculture (AFRI Grant No. 2012-67012-19883); to B.G.H., K.I.V., J.L.P., J.A., and T.K.O. by the NSF (Graduate Research Fellowships); and to N.K.W. by the National Geographic Society (9097–12), the NSF (DEB-1256758), the John Templeton Foundation (#41855), and the National Institute of General Medical Sciences of the NIH (R35GM119816).

