## Supplementary Note and Figures for "Evolution of herbivory remodels a *Drosophila* genome"

To assist in validation of gene models, an independent long read assembly was generated from a partially inbred *S. flava* colony as follows:

Dovetail HiC library preparation and sequencing. *S. flava* flies used for the genome assembly were collected from a laboratory colony. The colony was founded from >150 larvae collected near Dover, NH, USA, and subsequently maintained for several years in the laboratory. 300 male flies were flash frozen and stored at -80°C. A Dovetail HiC library was prepared in a similar manner as described previously (Erez Lieberman-Aiden et al., 2009). Briefly, for each library, chromatin was fixed in place with formaldehyde in the nucleus and then extracted. Fixed chromatin was digested with DpnII, the 5’ overhangs filled in with biotinylated nucleotides, and free blunt ends were ligated. After ligation, crosslinks were reversed and the DNA purified from protein. Purified DNA was treated to remove biotin that was not internal to ligated fragments. DNA was then sheared to ~350 bp mean fragment size, and sequencing libraries were generated using NEBNext Ultra enzymes and Illumina-compatible adapters. Biotin-containing fragments were isolated using streptavidin beads before PCR enrichment of each library. The libraries were sequenced on an Illumina HiSeqX to produce 380 million 2x150 bp paired end reads.

PacBio Library and Sequencing. DNA samples were quantified using Qubit 2.0 Fluorometer (Life Technologies, Carlsbad, CA, USA). SMRTbell libraries (~20kb) for PacBio Sequel were constructed using SMRTbell Template Prep Kit 1.0 (PacBio, Menlo Park, CA, USA) using the manufacturer's recommended protocol. The pooled library was bound to polymerase using the Sequel Binding Kit 2.0 (PacBio) and loaded onto PacBio Sequel using the MagBead Kit V2 (PacBio). Sequencing was performed on two PacBio Sequel SMRT cells, using Instrument Control Software Version 5.0.0.6235, Primary analysis software Version 5.0.0.6236 and SMRT Link Version 5.0.0.6792.

Falcon Assembly. The genome assemblies were performed using the FALCON 1.8.8 pipeline from Pacific Bioscience. First, 70-fold whole-genome, single-molecule, real-time sequencing (SMRT) data of Scaptomyza flava was used as input to the traditional FALCON pipeline using a length cut-off that correspond to 50x coverage of data during the initial error-correcting stage. This resulted in 0.511 million error corrected reads with an N50 read length equal to 19.6 kb. Second, the error-corrected reads were processed by the overlap portion of the FALCON pipeline. The aligned reads were assembled in the third stage of FALCON into 3,561 primary contigs containing 443.7 Mbp with an NG50 contig length of 543.7 kbp. Finally, the assembly was polished through PacBio’s Arrow algorithm from SMRT Link 5.0.1, using the original raw-reads.

Scaffolding the assembly with HiRise. The input de novo assembly, shotgun reads, and Dovetail HiC library reads were used as input data for HiRise, a software pipeline designed specifically for using proximity ligation data to scaffold genome assemblies (Putnam et al, 2016). Shotgun and Dovetail HiC library sequences were aligned to the draft input assembly using a modified SNAP read mapper (http://snap.cs.berkeley.edu). The separations of Dovetail HiC read pairs mapped within draft scaffolds were analyzed by HiRise to produce a likelihood model for genomic distance between read pairs, and the model was used to identify and break putative misjoins, to score prospective joins, and make joins above a threshold. After scaffolding, shotgun sequences were used to close gaps between contigs.

Assembly metrics. The genome assembly of *S. flava* was scaffolded into 2,151 scaffolds covering 443.89 Mbp (N50 = 60.816 Mb) with a maximum length of 99.1 Mbp and with 1,418 gaps (0.03%).

Data availability. A publication on this assembly is in preparation, including an in-depth investigation of its strengths and limitations, so the data is not yet publicly deposited. The reads, assembly, and other data are available from the authors upon request.

References.

1. Putnam NH, O'Connell BL, Stites JC, Rice BJ, Blanchette M, Calef R, Troll CJ, Fields A, Hartley PD, Sugnet CW, Haussler D, Rokhsar DS, Green RE. Chromosome-scale shotgun assembly using an in vitro method for long-range linkage. *Genome Research*. 2016; 26: 342-50.

2. Lieberman-Aiden E, van Berkum NL, Williams L, Imakaev M, Ragoczy T, Telling A, Amit I, Lajoie BR, Sabo PJ, Dorschner MO, Sandstrom R, Bernstein R, Bender MA, Groudine M, Gnirke A, Stamatoyannopoulos J, Mirny LA, Lander ES, Dekker J. Comprehensive Mapping of Long-Range Interactions Reveals Folding Principles of the Human Genome. *Science*. 2009; 326: 289-293.

### SUPPLEMENTARY TABLES

**Table S1.** Trophic enrichment of $^{15}N$ isotopes in herbivorous and detritivorous insects.
**Table S2.** Genome assembly statistics.
**Table S3.** Genome benchmarking with core dipteran genes.
**Table S4.** Gene annotation benchmarking with core dipteran genes.
**Table S5.** Kinetic efficiency for mustard oils (PEITC) from GSTs in *S. flava*, other animals, and fungi.
**Table S6.** Repetitive element content in *S. flava* falls within the range across other *Drosophila* genomes.

### SUPPLEMENTARY FIGURES

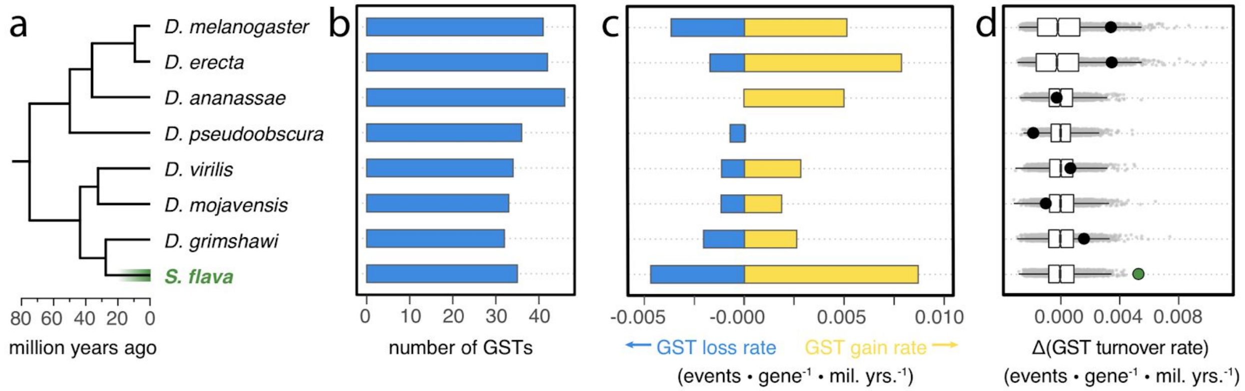


**Figure S1. Accelerated gain, loss, and turnover rates of GSTs in *S. flava*.** Results described in Fig. 3 of the main text are shown for GSTs in each taxon individually. Specifically, **(a)** The phylogenetic relationship for the eight focal taxa, with the dated phylogeny taken from (23). **(b)** Total GST number in each lineage. **(c)** Rates of GST gain and loss in each lineage. **(d)** Point estimates of branch-specific GST turnover rate (large circles) compared to turnover rates inferred for the same branches from a null distribution of 2,000 pseudo-datasets, simulated under the assumption of no branch-to-branch variation in turnover rate. The higher value observed in *S. flava* than in corresponding simulated datasets indicates the elevated turnover rate is statistically significant.

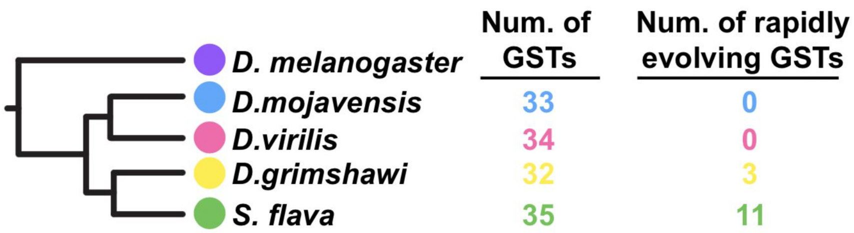


**Figure S2. Accelerated amino acid evolution of GSTs in *S. flava*.** The number of rapidly evolving GSTs in each lineage in the subgenus *Drosophila* was determined using branch tests for accelerated *dN/dS*. Significance was determined through likelihood ratio tests corrected for the false discovery rate ($q < 0.05$) within each lineage.

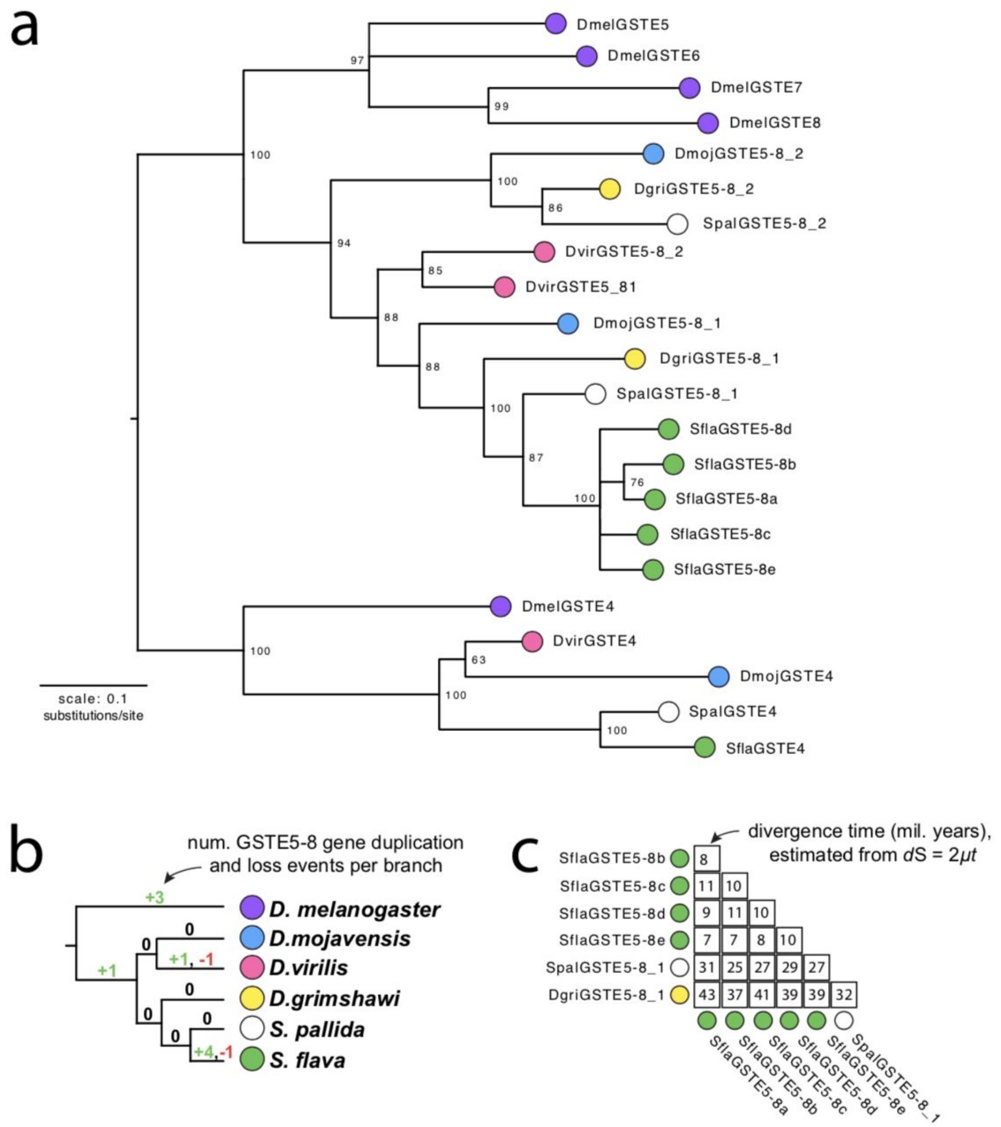


**Figure S3. The lineage-specific expansion of GSTE5-8 genes in S. flava occurred after the divergence from *S. pallida*, concurrent with the evolutionary transition to herbivory. (a)** Maximum likelihood nucleotide phylogeny of *GstE4* and *GstE5-8* orthologs identified in five of the species for which all *Gsts* were manually curated and in a fragmented short-read assembly of *S. pallida* (not yet published). Branches with bootstrap support values < 50 were collapsed. **(b)** *GstE5-8* gene duplication and loss events inferred by reconciling the phylogeny in panel "a" against the known species tree. **(c)** Estimated divergence times among *S. flava GstE5-8* paralogs and the most closely related *GstE5-8* ortholog from S. pallida and D. grimshawi. Divergence time was estimated using a Drosophila-specific mutation rate (104) and maximum likelihood estimates of pairwise synonymous site divergence, corrected for multiple substitutions.

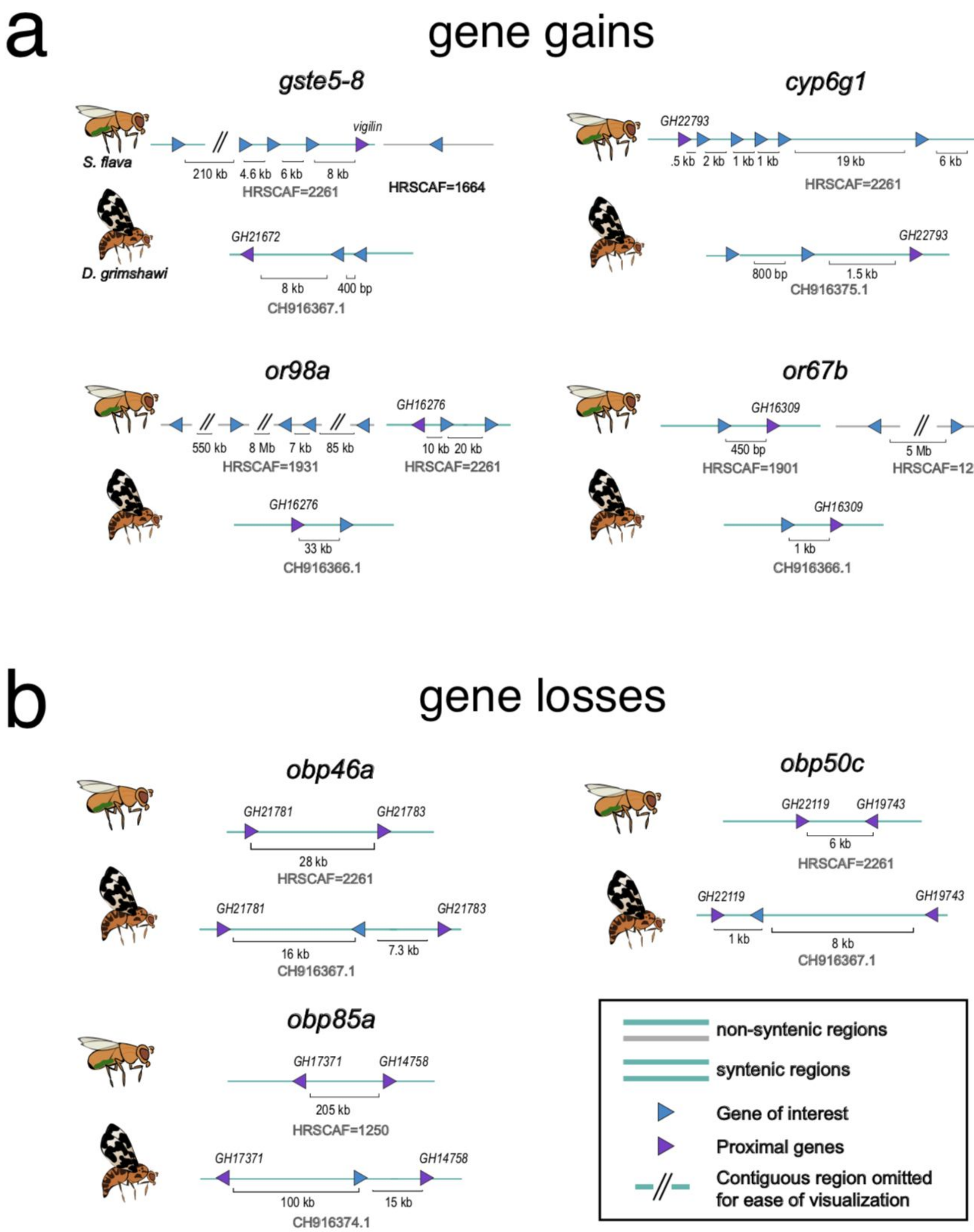


**Figure S4. Maps of the genomic locations of key gene gains (a) and losses (b) in *S. flava*.** These events were identified in the short-read assembly (ALLPATHS-LG + GapCloser). They were subsequently confirmed in the independent long-read assembly (HiRise) and, for gene gains, further confirmed by read depth analysis (see Fig. S6). Regions containing these scaffolds are shown for both *S. flava* and its closest non-herbivorous relative with a high-quality genome assembly, *D. grimshawi*. The "gene of interest" is indicated by name above above each plot. Scaffold names are indicated below; for *S. flava*, we depict scaffolds from the long-read (HiRise) assembly because it is less fragmented.

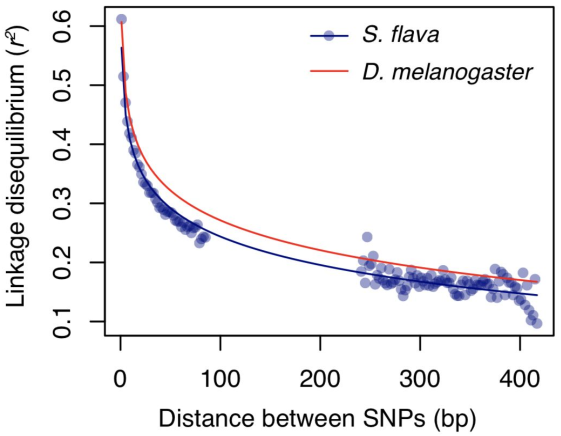


**Figure S5. Similarly rapid decay of linkage disequilibrium in *S. flava* and *D. melanogaster*.** LD for the fifteen largest autosomal scaffolds was estimated among pairs of sites that co-occur within paired-end Illumina reads from pooled sequencing of a wild-collected *S. flava* population. Points for *S. flava* represent average $r^2$ for SNP pairs binned by physical distance. The LD decay curve estimated using the same approach in *D. melanogaster* was taken from [110].

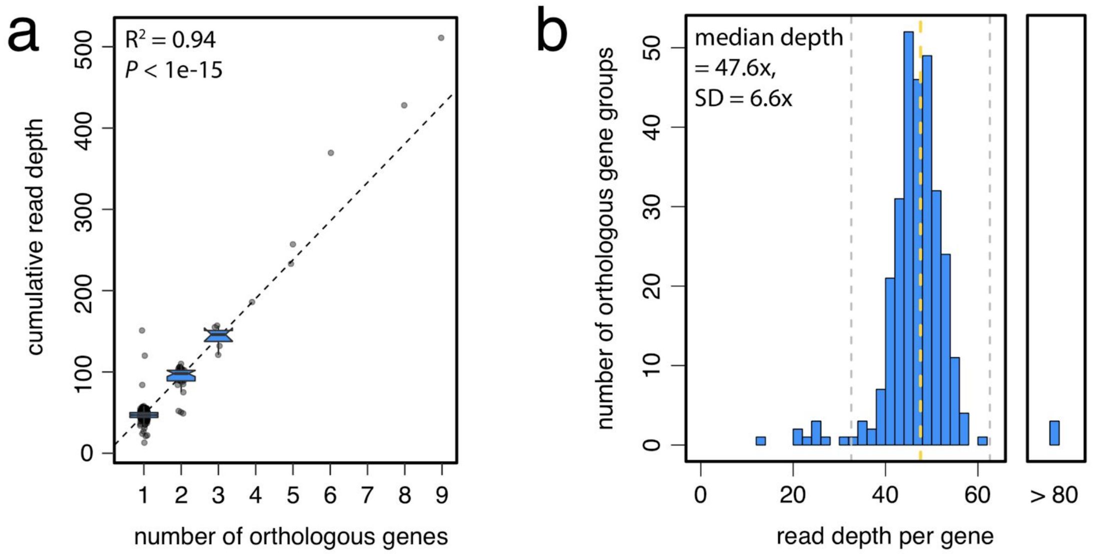


**Figure S6. Consistency between gene counts and short-read sequencing depth. (a)** Cumulative read depth per gene cluster is predictive of the number of genes annotated to that cluster. Note the narrow height of boxplots, signifying a tight density of points around the dotted line, whose slope is equal to the genome-wide median read depth after excluding poorly covered sites (depth < 5x). **(b)** For most genes, read depth per gene falls very close to the genome-wide median (dotted line), and the small number of outliers (deviations greater than the dashed lines in panel "b") were verified by agreement between both independent genome assemblies. Outliers with high read depth may be very recently duplicated gene copies (e.g., tandemly arrayed gene amplifications) that are too similar to assemble into separate genes, and/or contain repetitive sequences that may have collapsed during assembly; this is a general limitation of genome assemblies, but also appears to be uncommon here. This analysis was performed on the short-read genome assembly (ALLPATHS-LG + GapCloser).
